# Frequency of quorum sensing mutations in *Pseudomonas aeruginosa* strains isolated from different environments

**DOI:** 10.1101/2021.02.22.432365

**Authors:** Kathleen O’Connor, Conan Y. Zhao, Madeline Mei, Stephen P. Diggle

**Affiliations:** Center for Microbial Dynamics & Infection, School of Biological Sciences, Georgia Institute of Technology, Atlanta, GA 30332

**Keywords:** *Pseudomonas aeruginosa*, quorum sensing, LasR, cystic fibrosis

## Abstract

*Pseudomonas aeruginosa* uses quorum sensing (QS) to coordinate the expression of multiple genes necessary for establishing and maintaining infection. It has previously been shown that *lasR* QS mutations frequently arise in cystic fibrosis (CF) lung infections, however, there has been far less emphasis on determining whether other QS system mutations arise during infection or in other environments. To test this, we utilized 852 publicly available sequenced *P. aeruginosa* genomes from the *Pseudomonas* International Consortium Database (IPCD) to study *P. aeruginosa* QS mutational signatures. To study isolates by source, we focused on a subset of 654 isolates collected from CF, wounds, and non-infection environmental isolates, where we could clearly identify their source. We also worked a small collection of isolates *in vitro* to determine the impact of *lasR* and *pqs* mutations on isolate phenotypes. We found that *lasR* mutations are common across all environments and are not specific to infection nor a particular infection type. We also found that the *pqs* system proteins PqsA, PqsH, PqsL and MexT, a protein of increasing importance to the QS field, are highly variable. Conversely, RsaL, a negative transcriptional regulator of the *las* system, was found to be highly conserved, suggesting selective pressure to repress *las* system activity. Overall, our findings suggest that QS mutations in *P. aeruginosa* are common and not limited to the *las* system; however, LasR is unique in the frequency of putative loss-of-function mutations.

## Introduction

*Pseudomonas aeruginosa* is a Gram-negative opportunistic pathogen that can be problematic in a number of infection types and environments [1]. One of the major adaptations of *P. aeruginosa* during chronic infection is the loss of quorum sensing (QS) [2–8]. In *P. aeruginosa*, QS regulates the expression of hundreds of genes, including those that encode for secreted products and virulence factors [9, 10]. It is regulated via a complex hierarchical network, composed of two *N*-acyl homoserine lactone (AHL) circuits known as LasR-LasI and RhlR-RhlI, two orphan regulators termed QscR and VqsR, and a negative transcriptional regulator of the *las* system, RsaL [9–19]. The Las and Rhl systems are composed of LuxR-LuxI pairs, which are homologous to other Gram-negative bacterial QS systems. The LuxR-type receptors (LasR, RhlR) act as transcriptional regulators, and the LuxI-type proteins (LasI, RhlI) are signal synthases. LasI produces 3-oxo-dodecanoyl-L-homoserine lactone (3OC12-HSL), and RhlI produces N-butanoyl-L-homoserine lactone (C4-HSL). Both signals can function in a combinatorial manner to synergistically regulate genes [20, 21]. Working in conjunction with the two AHL systems, is an alkyl-quinolone (AQ) system, comprising the *pqsABCDE* operon and *pqsH, pqsL* and *pqsR* (*mvfR*) genes. These genes drive the synthesis and response of 2-heptyl-3-hydroxy-4-quinolone (the *Pseudomonas* quinolone signal; PQS) [22, 23].

LasR was first identified in 1991 as a regulator of the *lasB* (elastase) gene [24]. It has since been described as a key QS regulator in the well-studied laboratory strains PAO1 and PA14, where it has been shown to sit at the top of the QS hierarchy, regulating both the *rhl* and *pqs* systems [9, 11–15]. *lasR* mutants have frequently been isolated from cystic fibrosis (CF) lungs and more recently, it has been shown that some CF strains use RhlR to regulate the *rhl* and *pqs* systems in the absence of functional LasR [25–30]. The decoupling of the AHL QS hierarchy reportedly requires the inactivation of MexT [25, 27, 31], a regulator of the multi-drug efflux pump operon MexEF-OprN [32, 33]. It has also been shown that *mexT* mutation in PAO1 can decouple public *rhlR-*regulated traits from private metabolic *lasR-*regulated traits [20]. PqsE and RhlR have also been suggested to function as a ligand:receptor pair in some QS ‘re-wired’ strains [34–36].

Previous work has shown that *lasR* mutations are also found outside the CF lung, but the extent and variation of these mutations is still being elucidated [26, 30]. The degree of mutation in other QS genes in infection and environmental strains of *P. aeruginosa* remains unknown. In this study, we explored the diversity and frequency of QS mutations across a range of ecologically distinct environments to determine (i) which QS genes are frequently mutated; (ii) mutational signatures, or patterns in other genes associated with *lasR* mutations; (iii) *lasR* gene mutation frequency specific to isolate source; and (iv) the phenotypic outcome of QS mutations.

## Results

We utilized the published sequences of 852 *P. aeruginosa* isolates from the International *Pseudomonas* Consortium Database (IPCD); a database representing a range of *P. aeruginosa* strains from different sources including rivers, human infection and plants [37]. We queried key QS genes from the *las, rhl and pqs* systems, as well as the QS regulators *qscR, rsaL* and *vqsR* against gene sequences from PAO1 (GCF_000006765.1) for all 852 isolates. For some analyses, we additionally looked at *mexT* and *psdR*, two genes that have been associated with *lasR* mutations *in vitro* [25, 31, 38, 39]. We used two analysis pipelines to look at the variation in QS genes in *P. aeruginosa*. One pipeline used NCBI BLASTn [40] and BLOSSUM80 [41], and the second analysis utilized GAMMA [42]. All analyses were conducted in R version 4.3 [43].

### Quorum sensing genes are highly variable at the nucleotide level

We first analyzed the diversity of nucleotide sequence for each QS gene for all 852 isolates and looked at which isolates had mutations in multiple QS-related genes. We found a large diversity in sequences and that few isolates had QS genes identical to PAO1 (GCF_000006765.1). The *rhl* system was especially variable, with 849 and 847 isolates differing from PAO1 for *rhlR* and *rhlI* respectively (Fig. 1A). The *las* system was more conserved, with 553 and 290 isolates varying from PAO1 for *lasR* and *lasI* respectively (Fig. 1A). Genes involved in 2-alkyl-4-quinolone (AQ) biosynthesis and response showed a large variation in mutations between genes, with *pqsE* as the least variable (479 mutated isolates), and *pqsH* and *pqsL* as the most variable (845, 847 respectively) (Fig. 1C). The orphan QS transcriptional regulators *qscR* and *vqsR* were highly variable (772 and 559 respectively). However, the negative *las* system transcriptional regulator *rsaL* was relatively conserved (218 mutated isolates) (Fig. 1D).

**Figure 1.**
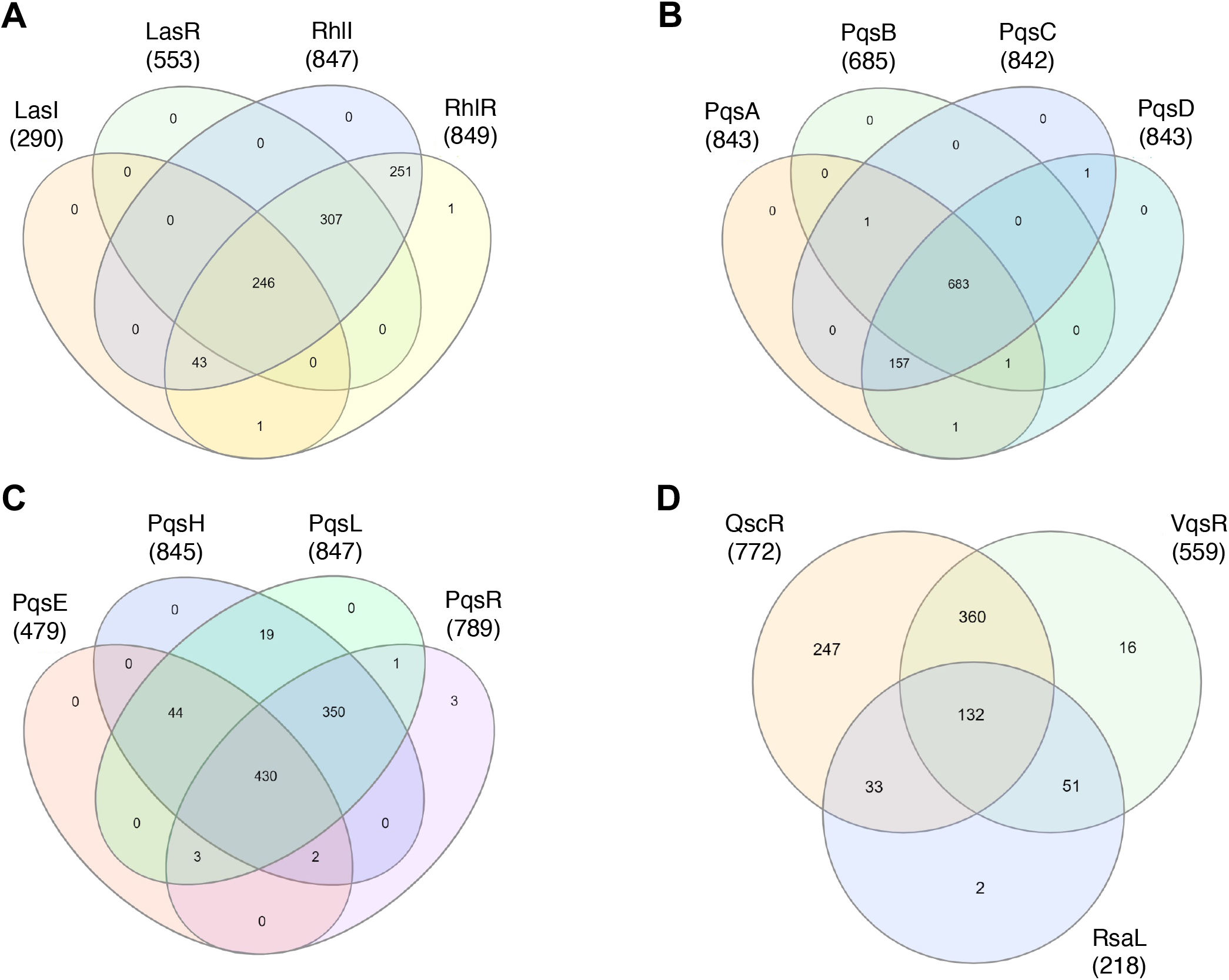
Comparison of nucleotide sequence variation among QS genes of *P. aeruginosa* isolates of the IPCD. There is a high degree of variation in nucleotide sequences in many of the QS genes, obscuring patterns in mutation overlap between genes within an isolate. (A) A Venn diagram of *las* and *rhl* system genes are plotted together, showing how many isolates share mutations in multiple genes. 246 isolates are mutated for all 4 genes, and few isolates have mutations in only a single gene. (B) A Venn diagram for *pqsABCD* shows a similar pattern and 683 isolates are mutated in all 4 genes, while few isolates are mutated in only one *pqs* gene. (C) Few isolates have only one *pqs* gene mutation, and 430 isolates have mutations in all *pqsEHLR* genes. (D) A Venn diagram of the negative QS regulators *vqsR, rsaL, and qscR* show that 132 isolates share mutations in all 3 genes.

### Negative and positive regulators of the *las* system are antipodal in sequence variation

We counted the number of unique protein sequences found in our database for each QS gene. We added *mucA* to our analysis as a reference gene that is frequently mutated in CF lung infections [44, 45], and *rpsL*, which encodes the 30S ribosomal protein S12; a conserved housekeeping gene [46, 47]. When we queried each QS gene nucleotide sequence against the 852 isolates using BLASTn, the query returned less than 852 sequences for each gene. This disparity is likely due to gaps in sequences, gene deletions, and extensive mutations, preventing BLASTn from returning a query. Additionally, for this analysis, we removed all truncated nucleotide sequences from the analysis, to avoid including sequencing near the end of a contig. Fig. 2A shows the number of protein sequences analyzed for each gene. After MucA, a LasR query returned the fewest number of sequences – 756 out of 852 isolates, suggesting that there are many strains that contain large deletions in LasR, truncations, or are lacking the LasR gene entirely. After translating the sequences, we found that LasR had the most unique protein sequences (259) across 852 isolates (Table 1, Fig. 2A) compared to other QS genes and MucA. The next most variable QS proteins, PqsH, PqsA, and PqsL contained 189, 170, and 169 unique protein sequences, respectively (Table 1; Fig. 2A). We found that AHL signal synthases were highly conserved; LasI and RhlI had only 61 and 87 unique protein sequences respectively. RsaL was the most conserved of all studied proteins, with only 18 unique sequences. Compared to LasR, the other key QS proteins were more conserved across isolates. Our protein sequence findings appear to contrast with our nucleotide sequence analysis, but this is likely due to primarily silent mutations in the *rhl* system. To control for possible bias due to differences in protein length, we normalized gene sizes to PAO1 by dividing the lengths of each gene by the PAO1 reference gene length. We found that the shortest proteins, RspL and RsaL were the most conserved, while the longest genes, PqsH, PqsA, and PqsL were all highly variable. The most variable protein, LasR is of intermediate length, compared to RsaL and Pqs proteins (Table 1).

**Table 1.** Variation in protein sequence encoded by *P. aeruginosa* isolates. Using the sequences returned by BLASTn, we translated genes with full length nucleotide sequences to determine the number of unique protein sequences encoded by the isolates in the database. We found that LasR had the most unique sequences across queried isolates, and RsaL encoded the fewest unique sequences.

| Table 1. Protein Data |  |  |  |  |
| --- | --- | --- | --- | --- |
| Gene | Gene Length | Genes Returned | Unique Protein Sequences | Unique Sequences/<br>Gene Length |
| *rsaL* | 243 | 806 | 18 | 0.07 |
| *rpsL* | 372 | 829 | 27 | 0.07 |
| *lasI* | 606 | 809 | 61 | 0.10 |
| *rhII* | 606 | 827 | 87 | 0.14 |
| *vqsR* | 807 | 816 | 89 | 0.11 |
| *qscR* | 714 | 823 | 98 | 0.14 |
| *pqsB* | 852 | 822 | 103 | 0.12 |
| *pqsE* | 906 | 818 | 117 | 0.13 |
| *mucA* | 585 | 713 | 120 | 0.21 |
| *pqsR* | 999 | 818 | 122 | 0.12 |
| *pqsD* | 1014 | 820 | 125 | 0.12 |
| *pqsC* | 1047 | 823 | 128 | 0.12 |
| *rhIR* | 726 | 819 | 133 | 0.18 |
| *pqsL* | 1197 | 827 | 169 | 0.14 |
| *pqsA* | 1554 | 790 | 170 | 0.11 |
| *pqsH* | 1149 | 829 | 189 | 0.16 |
| *lasR* | 720 | 756 | 259 | 0.36 |

**Figure 2.**
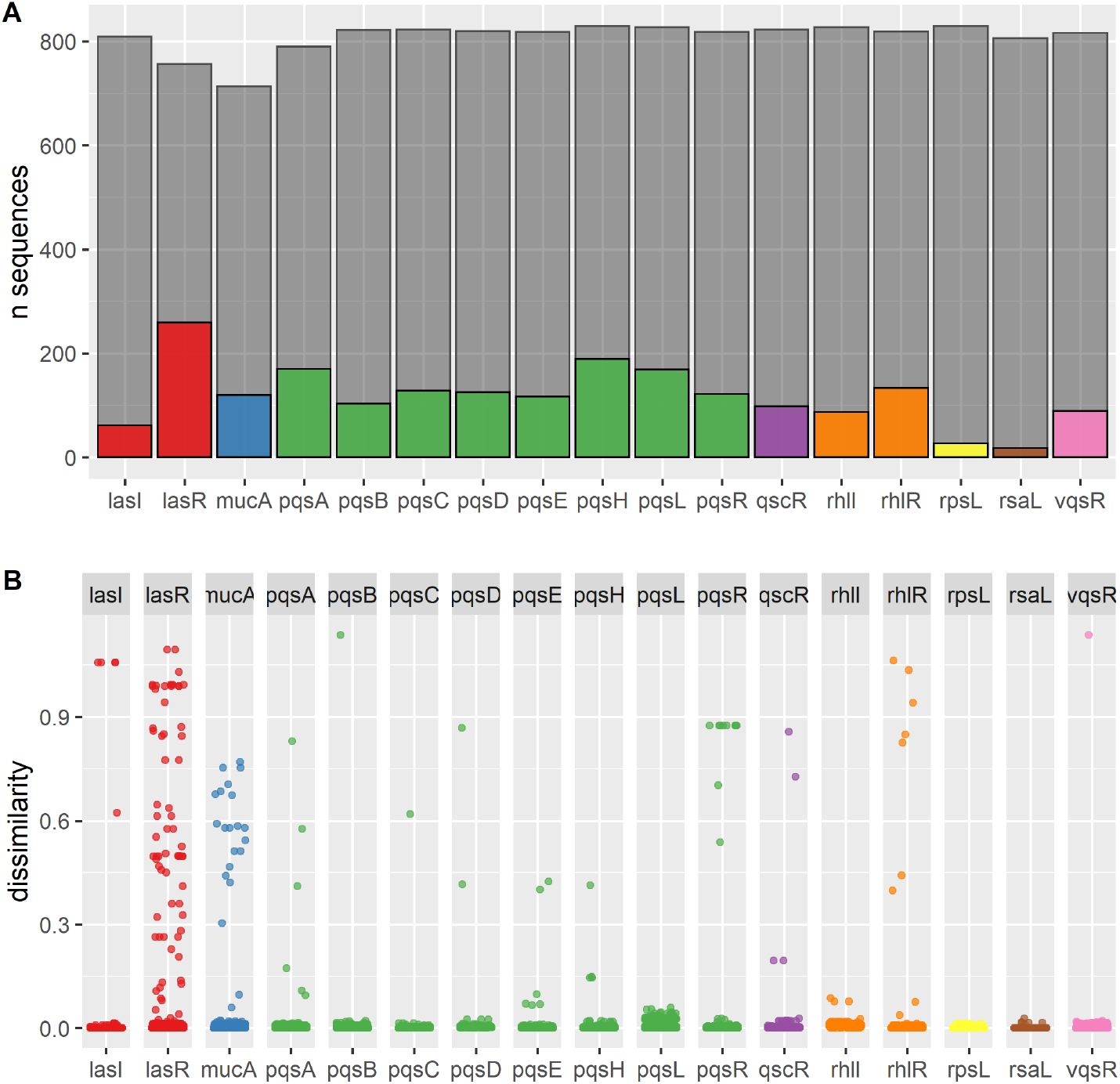
Determining variability in QS proteins between *P. aeruginosa* isolates from the IPCD. (A) We created a database of 852 isolates and used PAO1 to search for the QS proteins of each isolate. Due to the variation in each isolate’s genome and due to gaps in sequencing, each protein queried returned fewer than 852 sequences (shown in gray). We also determined the number of unique sequences for each protein and found that LasR had the highest number of unique sequences (in color). The next highest were PqsH, PqsA, and PqsL. RsaL had the fewest number of unique sequences. (B) Using a custom dissimilarity metric (BLOSUM80), we calculated mean dissimilarity scores. We found that LasR had the highest mean dissimilarity score compared to all QS proteins, and the largest variation.

We next plotted the BLOSUM80 dissimilarity scores for all isolates for each gene and found that LasR had a large number of highly dissimilar sequences (Fig. 2B). The highly dissimilar scores are caused by truncation mutations due to an early stop site leading to a shortened protein. Transcriptional regulators PqsR and RhlR also have a number of truncation mutations, however, there are fewer than LasR. While PqsA, PqsL and PqsH all encode many unique protein sequences, most isolates cluster towards the bottom, indicating that they have dissimilarity scores close to 0, indicating that the proteins have a near identical function.

### MexT, PqsL and LasR have the highest mutation rates

We used the protein sequence data from GAMMA to look at the mutation rate of each QS protein (Table 2). For this analysis, we were able to include truncated nucleotide sequences, and only excluded sequences where it was indicated that the gene was located at the contig edge. We included MexT and PsdR proteins in this analysis. For the majority of genes, we compared isolates to the PAO1 (GCF_000006765.1) gene sequence, with the exception of PqsD and RhlI which we compared to the PAK strain of *P. aeruginosa*. The MexT sequence for PAO1 has an 8 bp insertion sequence, which was not commonly found in the IPCD database of isolates, as well as many other coding mutations. We instead used the sequence for isolate U0330A as it coded for a protein shared by 128 other isolates, the highest number of strains sharing one MexT sequence. The MexT gene is highly variable, and is mutated in our Nottingham PAO1 lab strain (NPAO1) and the PAO1 reference used (GCF_000006765.1). Among the QS genes, we found that PqsL had the highest mutation rate, followed by LasR. We defined mutation rate in this analysis as the percentage of non-wildtype protein sequences out of queried isolates. RsaL and LasI were highly conserved, with low mutation rates.

**Table 2.**
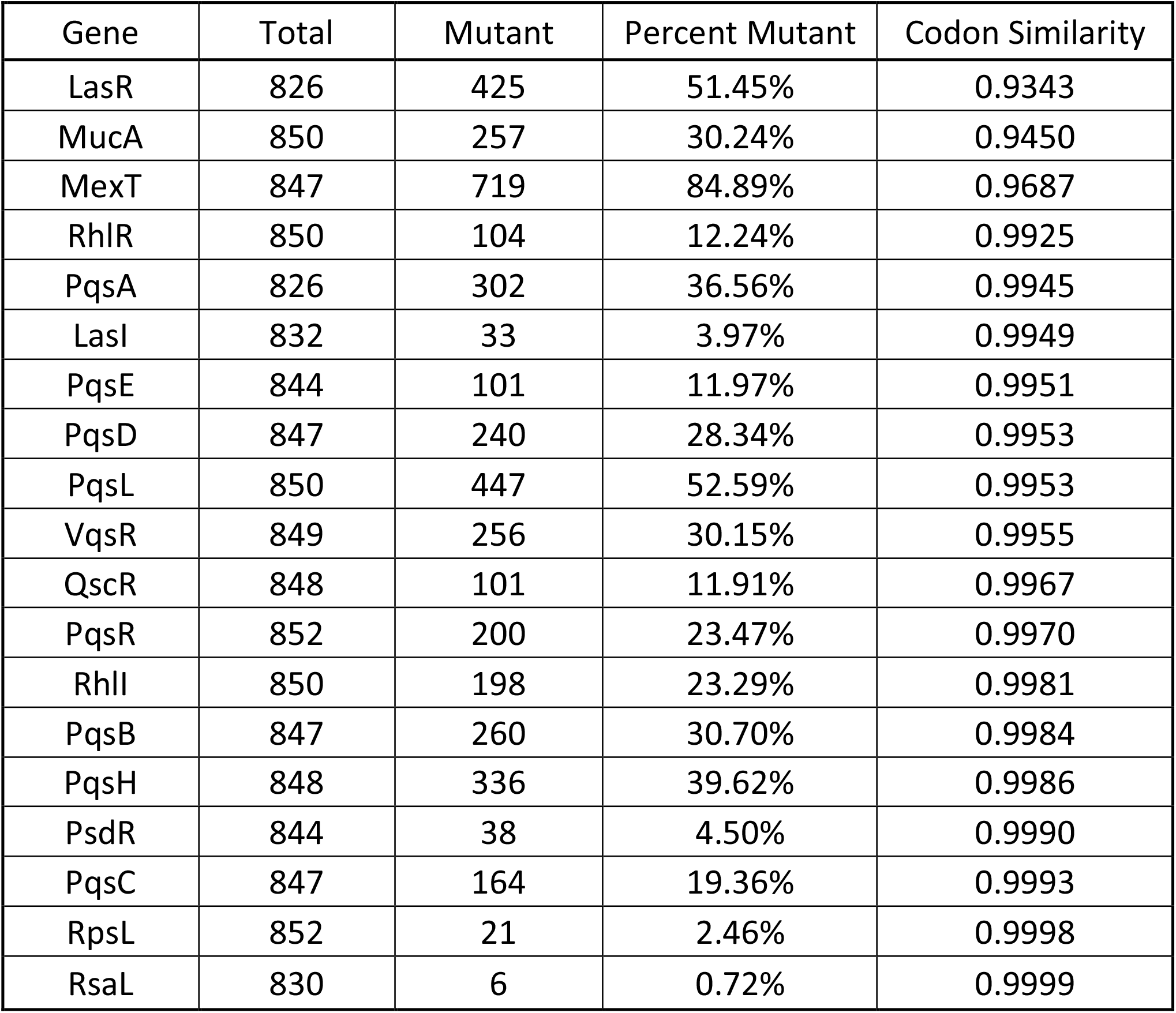
Mutation rate and mean codon similarity of QS proteins in *P. aeruginosa*. We used GAMMA to query QS genes against 852 isolates. We used PAK as a reference for RhlI and PqsD, U0330A as a reference for MexT and PAO1 as a reference for all other genes. GAMMA assigns isolates with a codon similarity measure which reflects the type of amino acid change or truncation. A low score reflects a likely change in function of the protein, and a score of 1 reflects the isolate is identical to the reference. We found that LasR had the lowest mean codon similarity and RsaL the highest.

Using GAMMA generated protein mutation data, we next calculated the average codon similarity of all isolates to a reference (Table 2). High codon similarity suggests similar protein function, while low codon similarity indicates a change or loss of protein function. For the majority of genes we used PAO1 (GCF_000006765.1) as a reference, however the codon similarity average for several proteins were low. To address this, for RhlI and PqsD we used PAK as the reference strain, and for MexT we used U0330A as these strains were likely to be better models based on higher similarity averages for each protein. Similar to our analysis using BLOSUM80, we found that LasR had the lowest average protein similarity of all QS genes, followed by MexT, which we attributed to the high rate of truncation mutations. All other QS proteins had a codon similarity average of greater than 0.99, indicating that while there is variation in the protein sequence, many of these mutations likely lead to a similarly functioning protein.

### LasR protein mutations are common across all environments but are more divergent in human wounds

In agreement with previous studies, LasR was the most variable gene in our protein analysis (Table 1; Fig. 2), and we found that the highly dissimilar LasR sequences were due to protein truncations [2, 30, 48]. To determine if LasR mutations vary by environment, we categorized the strains by source. Using data from the IPCD, we selected a subset of 654 strains labeled as “environmental”, “cystic fibrosis” or “CF”, and “wound” or “ulcer” or “burn” and reclassified them as environmental (209 strains), CF (396 strains), or wound (wound, ulcer, and burn) (49 strains); 654 total. The remaining 198 strains from the original set of 852 strains were of uncertain origin and therefore excluded from this analysis. To establish a threshold by which a protein could be deemed functional or not, we looked at truncated LasR proteins within each environment. We compared the amino acid length of LasR in the IPCD strains to the PAO1 LasR protein – which is equal in length to many commonly researched strains including PA14, PAK and the Liverpool epidemic CF strain LESB58. Our assumption was that a truncated protein due to an early stop site, would lead to a nonfunctional protein. We used a stringent 100% length as a cutoff, and any protein shorter than full-length was considered truncated. Fig. 3A shows the proportion of each group that had truncated LasR proteins with CF, environmental, and wound isolates having 20%, 11% and 30% truncations respectively. We then plotted the protein sequence dissimilarities categorized by isolate source, and found highly dissimilar isolates, primarily in CF and wounds (Fig. 3B). Overall, we found that *lasR* mutations are ubiquitous across all environments, but there is a larger percentage of strains with truncated LasR proteins found in infection environments. We additionally calculated the mean and median codon similarity scores for each environment, with 1 being a protein identical to PAO1. We found that the wound group was the most dissimilar compared to PAO1, with the lowest mean similarity score (0.893), and CF and environmental were similar (CF: 0.953, Env: 0.948). The wound group also had the lowest median similarity score (0.995), followed by CF (0.996) and environmental had the highest median similarity score (1.0).

**Figure 3.**
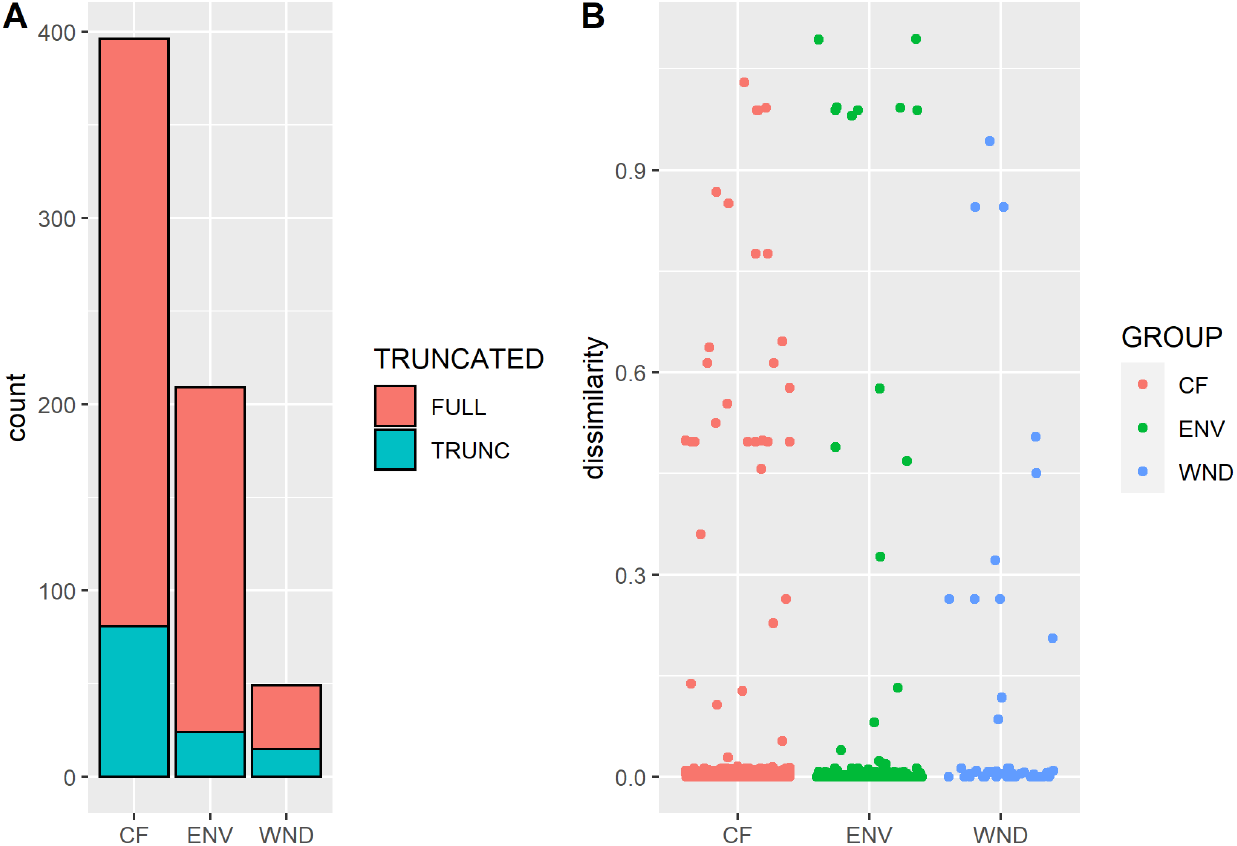
LasR truncations are found in isolates from all sources. To observe the fraction of truncated proteins across all environments, we categorized the isolates into 3 groups: cystic fibrosis (CF), environmental (ENV) or wound (WND). (A) We show the number of truncated proteins out of the total number of isolates in each group. The protein variation plot (B) depicts the similarity scores for all groups compared, and we see the highly dissimilar truncated proteins consisted primarily of CF and WND isolates.

### *lasR* mutation is not a predictor of exoprotease production or colony autolysis in selected environmental and infection isolates

To accompany our genomic analysis, we selected 12 isolates from the IPCD in order to perform phenotypic assays to assess QS function. We included our lab strain NPAO1 as well as the clean deletion mutants PAO1Δ*lasR*, PAO1Δ*rhlR*, PAO1Δ*lasR*Δ*rhlR*, PAO1Δ*lasI*. Our PAO1Δ*lasR* strain made high levels of protease that were not significantly lower than NPAO1 (89 % of NPAO1 levels, *p* = 0.338). A *lasR* gentamicin insertion mutant made 75 % of the protease levels of NPAO1 (*p* = 0.113) (Fig. 4). We found that unlike mutations in LasR, mutations in RhlR had a significant effect on the protease levels produced compared to NPAO1 (PAO1Δ*rhlR*: 45 %, *p* = 0.013; PAO1Δ*lasR*Δ*rhlR* double mutant: 20 %, *p* = 0.011). Compared with both Δ*lasR* mutants, NPAO1Δ*lasI* made lower amounts of protease (60 %, *p* = 0.025) (Fig. 4). Isolates with mutated LasR proteins that produce significantly less protease than wildtype included A17, CND03, CPHL2000, Jp115 (*p* values respectively: 0.01, 0.01, 0.02, 0.01). PT31M, a German environmental isolate contains a truncation mutation in LasR, yet it produced an intermediate amount of protease compared to NPAO1 with an intact LasR (69 %, *p* = 0.062). The Belgian river environmental strain W15Dec14 also produced an intermediate amount of protease (67 %, *p* = 0.065) and is wildtype for LasR, but does contain coding mutations in RhlI. All other isolates with wildtype LasR proteins produced protease levels that were not significantly different from NPAO1 (Fig. 4).

**Figure 4.**
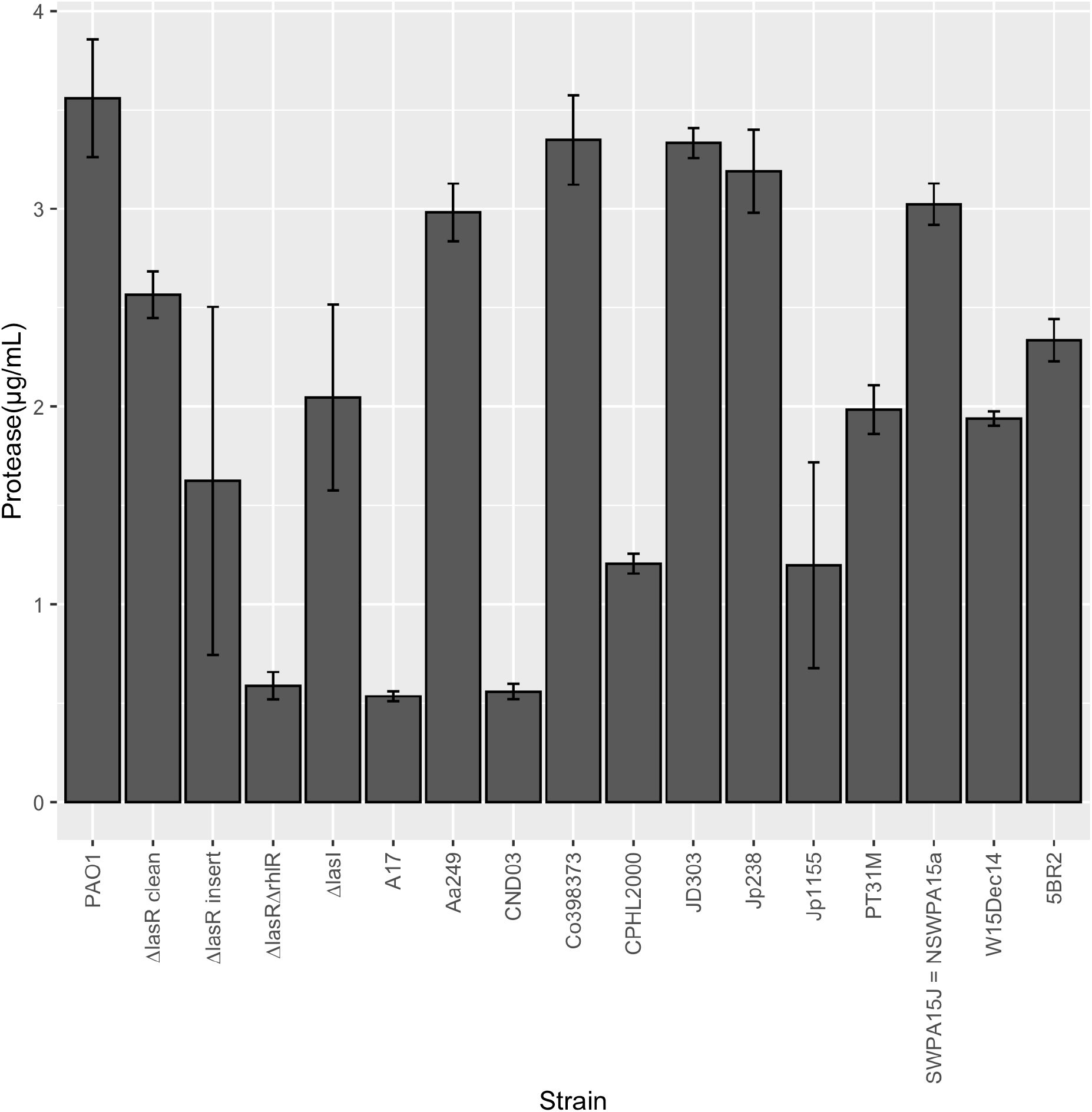
Exoprotease secretion by environmental and clinical isolates. We measured the supernatant exoprotease levels produced by stationary phase isolates, compared to PAO1, PAO1Δ*lasR* and PAO1Δ*lasR*Δ*rhlR*. We normalized the readings by optical density (OD 600_nm_) and compared all isolates and clean mutants to PAO1.

Lysis in QS mutants has long been observed in the lab, specifically in *lasR, lasI* and/or *pqs* mutants [48–51]. We used QS deletion mutant strains to determine how QS mediates colony autolysis. PAO1Δ*lasR* and PAO1Δ*pqsL* showed similar lysis phenotypes and a metallic sheen, while PAO1Δ*pqsH* caused colony plaquing (Fig. 5). PAO1Δ*pqsA*Δ*lasR*, PAO1Δ*pqsA*Δ*pqsH* and PAO1Δ*pqsA*Δ*pqsL* mutations led to a loss of colony autolysis (Fig. 5). We found that all LasR wildtype strains we tested from the IPCD showing lysis, contained multiple mutations in the *pqs* quinolone system, as follows: SWPA15J=NSWPA15a (PqsC,D,E), 5BR2 (PqsD,L), Co398373 (PqsD,L), Jp238 (PqsB,D,L), W15D14 (PqsE,D,L), Aa249 (PqsB,D,L), JD303 (PqsA,B,D).

**Figure 5.**
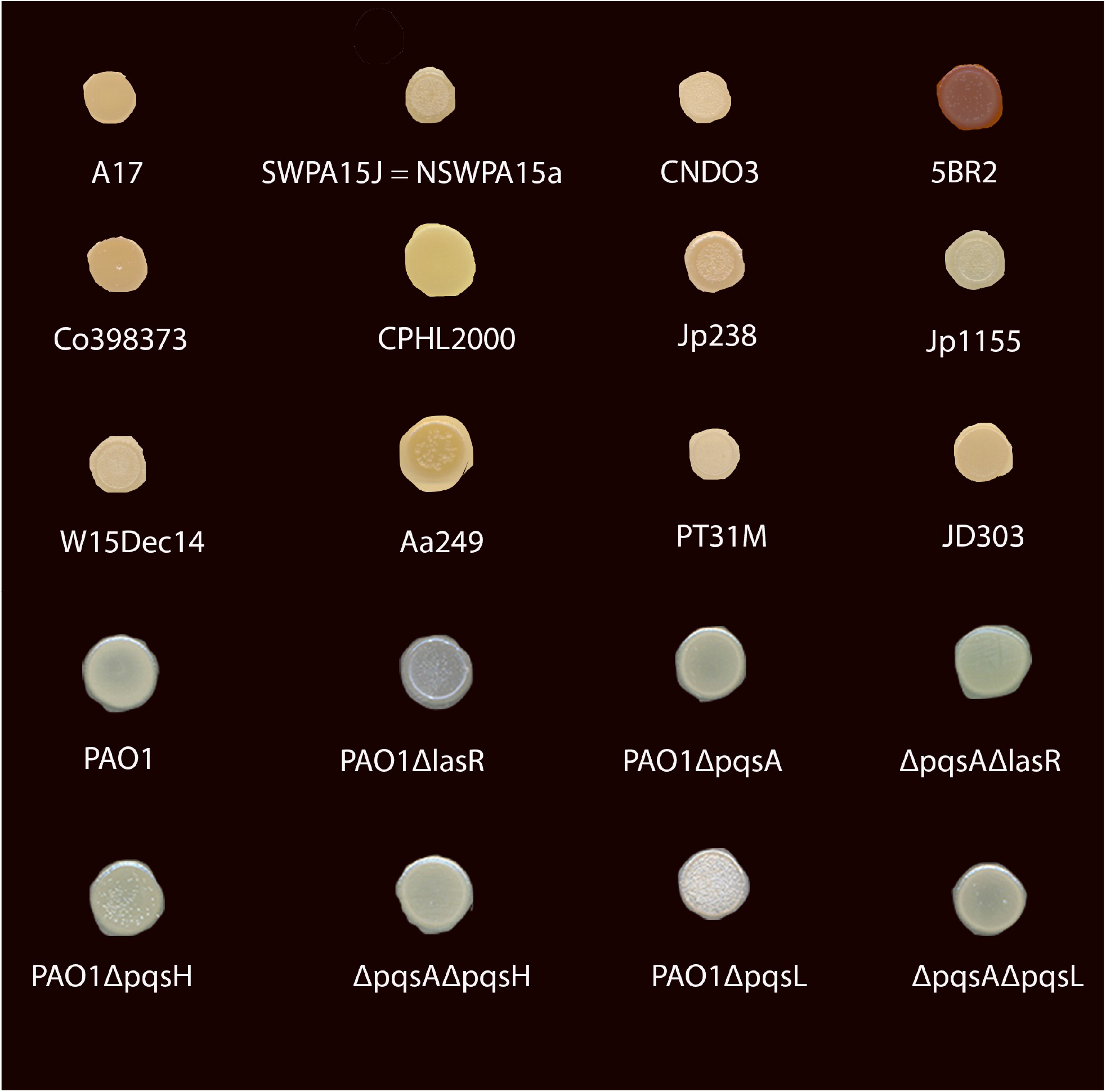
Colony autolysis phenotypes of isolates compared to lab strain QS mutants. We spotted overnight cultures on LBA plates to observe the colony autolysis phenotypes exhibited by clean QS deletion mutants alongside environmental and clinical isolates.

Strains with wildtype LasR showing lysis were from mixed environmental, wound and CF sources. There were two *lasR* mutant isolates that showed no lysis - A17 and CPHL2000, both sourced from human wounds (Fig. 5). Isolates A17 and CPHL2000 have 1 and 2 SNPs in LasR respectively, both creating a change in amino acid sequence (A17: L110P, CPHL2000: E196G). As expected, the 3 isolates with truncated LasR (CND03, Jp1155, and PT31M) demonstrated lysis.

## Discussion

In *P. aeruginosa, lasR* QS mutants are frequently isolated from human chronic infection and environmental sources, but less is known about other QS genes [2, 48, 52]. Using a publicly available database of 852 fully sequenced isolates from CF, wounds and non-human associated (environmental) strains [37], we determined the mutation frequency and variation of *lasR* and other QS genes in *P. aeruginosa*. To determine how QS genotype impacts phenotypic function, we used 12 strains that were sequenced as part of the IPCD study [37]. We found that (i) by multiple metrics, LasR is the most variable QS protein; (ii) LasR mutations are found in isolates across all environments, suggesting that any environment can drive the evolution of these mutations; (iii) the negative *las* system regulator RsaL is well-conserved; (iv) signal synthases LasI and RhlI are conserved compared to their transcriptional regulator pairs; (v) coding mutations in *pqs* system genes are common and may impact colony autolysis, however they are less divergent than LasR mutations; (vi) MexT is highly variable and mutations are common.

Our work supports the long-held belief that LasR is a commonly mutated QS protein as we showed that LasR has the highest number of unique protein sequences compared to other QS genes. A high number of isolates carried mutations in LasR compared to the PAO1 reference protein sequence. The IPCD strains also had the lowest average codon similarity for the LasR gene when compared to other QS regulon genes, which we attributed to nonsense mutations leading to protein truncation, and other large changes in amino acid sequence. Our results also highlight that LasR mutations are found in environmental isolates and are not specific to human infections. Truncation mutations are more commonly found in human wound infections, suggesting that there might be some selective pressure to evolve loss of function mutations. Mean and median similarity scores suggest there is not a large difference between CF and environmental isolates, and instead we found that wound isolates are enriched for loss of function mutations. The wound group was the smallest group of isolates we queried thus a larger exploration of wound strains could provide a deeper understanding of this phenomenon.

Interestingly, the most highly conserved QS protein was RsaL, which negatively regulates the *las* QS system [16, 53, 54]. RsaL and LasR are antipodal in variation across isolates and perform opposite jobs in regulating the *las* system. There may be a benefit to tightly conserving the “off” switch of the *las* system while frequently mutating and losing the function of the “on” switch, suggesting that a key element of the QS regulatory cascade is down-regulating *las*-controlled QS in *P. aeruginosa*. Given this complexity, it remains a challenge to understand why the *las* system evolved and what fitness benefits it provides *P. aeruginosa* in different environments. There may be evolutionary benefits for both the maintenance and loss of LasR so that both *lasR* positive and negative strains can stably coexist in heterogenous populations and contribute to an overall community function. In support of this idea, it has been shown that (i) *lasR-* strains overproduce Rhl-associated factors and cross-feed wild type cells in low iron environments, which will likely impact infection dynamics of mixed populations [55]; (ii) mixed *lasR* +/- populations display decreased virulence in mouse models of infection [56]; and (iii) mixed populations exhibit enhanced tolerance to beta-lactam antibiotics [57]. These data suggest there are likely considerable fitness advantages to cells growing in heterogeneous QS populations, perhaps as a bet-hedging mechanism for future disturbance events.

The signal synthases of the two AHL QS systems in *P. aeruginosa* are conserved compared to their transcriptional regulator pairs. It has been hypothesized that there could be evolutionary mechanisms to conserve signal synthases while mutating transcriptional regulators [58]. We saw a significant difference in protease production by the clean deletion of the regulator mutant LasR and the signal synthase mutant LasI, where deletion of LasI resulted in a larger decrease in protease production. The reason for this could be further explored and may be impacted by the MexT mutation in NPAO1, common to other PAO1 lab strains, or the fact that the 3’ end of RsaL is also deleted in a clean LasR deletion mutation [59]. There could also be an evolutionary benefit to maintaining some protease function in the absence of fully functioning QS, supporting the maintenance of signal synthases while mutating positive transcriptional regulators.

Recent studies on QS in *P. aeruginosa* has revealed that the complex and intertwined *las, rhl* and *pqs* systems can be re-wired if *lasR* becomes mutated [25, 27, 31, 34–36]. It is not always clear whether these strains are entirely QS-null or if they have re-wired their QS systems to circumvent the loss of *lasR*. In our *in vitro* work, we looked at lysis and protease secretion as indicators of QS function, which have previously been used to screen for LasR function [48, 60]. We found an environmental isolate, PT31M, with a truncated *lasR* that produces high levels of protease. PT31M may be an example of a re-wired isolate showing QS independent of LasR, suggesting this adaptation is not specific to CF lung infections. Colony lysis has been used as a predictor for *las* system function, however this study has shown little correlation between lysis and *lasR* genotype, potentially due to *pqs* mutation mediated lysis seen in NPAO1Δ*pqsH* and NPAO1Δ*pqsL*.

Overall, our work shows that LasR is uniquely mutated compared to other QS genes, future work should more strongly focus on the ecology of mixed QS-phenotypes to better understand QSinvolvement in infection and other environments. With ongoing work identifying QS-inhibitors targeting the *las* QS system, the frequency of *lasR* mutated strains found in our study suggests that this particular pursuit might be improved by targeting a less variable QS gene, such as *rhl* system genes or RsaL.

## Materials and Methods

### Querying QS genes from the International *Pseudomonas* Consortium Database

Using nucleotide sequences from PAO1 (GCF_000006765.1), we queried QS genes using BLASTn for isolates from the IPCD [37]. We chose this strain because it is a fully sequenced, frequently used lab strain. We first compared strains with nucleotide mutations relative to PAO1 in each of the QS genes of interest, to determine how frequently strains exhibit nucleotide polymorphisms across multiple QS genes. We then translated these sequences into protein sequences calculating putative amino acid sequence similarities using BLOSUM80 [41]. First, we compared genes found in each isolate against our reference strain, PAO1, normalized against the similarity of the reference against itself. We then calculated the mean dissimilarity score of all isolates compared to PAO1. Some isolates were missing genes due to sequencing errors or true truncations, the number of isolates with a given gene present was under 852 for all genes. All analyses, including translation steps were conducted in R version 4.3. All code and files are available on a publicly available database: https://github.gatech.edu/koconnor36/Frequency_of_quorum_sensing_mutations_in_Pa2021

### Creating an IPCD database using BLASTn

We pulled IPCD data from GenBank from the PRJNA325248 BioProject (https://www.ncbi.nlm.nih.gov/bioproject/325248), and downloaded contigs as a multifasta file. We used the makeblastdb/ command to generate a database of all isolate contigs.

### Using BLASTn to find QS genes for each isolate

Using our generated database, we queried the PAO1 sequence for each gene found on Pseudomonas.com [61], against the database. We generated csv files for each gene which included the gene sequences for each isolate.

### Comparing strains with nucleotide polymorphisms in multiple QS genes

We used the csv files for each gene generated from the BLASTn analysis to isolate all accession information of strains with <100% query coverage and <100% identity to the PAO1 sequence of each QS gene. For each QS gene, we used this list of NCBI accession IDs, corresponding to unique strains, to visualize the number of strains with nucleotide mutations in one or more genes with InteractiVenn [62]. We then calculated the number of strains and the percentage of strains in the IPCD database with at least one mutation in each QS gene. We generated accession lists of mutated strains for each QS gene using a custom R script (v.4.0.2) and generated Fig. 1 with InteractiVenn. All analyses were conducted in R version 4.3.

### Translating nucleotide to amino acid sequence

We translated genes to proteins using a custom R script. We first queried only for sequences starting with a canonical ATG start codon. VqsR is an exception as it begins with the alternative start codon “GTG”. We translated the sequences meeting these criteria using the translate function from the BioStrings R package (v.2.58.0) [63].

### Calculating dissimilarity scores for isolates’ QS proteins

All sequence analyses were performed in R (v.4.0.2) using the Biostrings package v.2.58.0. We compared isolate protein sequences to PAO1 protein sequences using BLOSUM80, a matrix designed to compare protein sequences within species.

### Determining truncation rates for LasR and categorizing isolates by location

We determined the length of the reference LasR protein, from PAO1, compared to each isolate protein. If the isolate protein was 100% or less of the length of the PAO1 protein, we categorized it as truncated. Sequences were categorized as CF-originated (CF), environmental (ENV), or wound (WND). If the sequence was entered into IPCD as environmental, we adopted that label. Additionally, we included sequences labeled from animal hosts as environmental. For CF, we only included sequences with sources explicitly labeled as CF or cystic fibrosis. For wound, we included sequences labeled as wound, ulcer, and burn.

### Protein mutation analysis using GAMMA

We used GAMMA [42] to find the mutations in QS genes for all isolates. We created a multifasta of genes: *lasR,I, rhlR,I, pqsA,B,C,D,E,H,L,R, qscR, rsaL, vqsR, mucA, rpsL, mexT, psdR* to query against the IPCD PRJNA325248 multifasta. We ran commands as described in the GAMMA GitHub [42] https://github.com/rastanton/GAMMA, and generated a GAMMA file with mutations reported for all genes in all IPCD isolates. All other sequences are available on our project Github https://github.gatech.edu/koconnor36/Frequency_of_quorum_sensing_mutations_in_Pa2021.

### Phenotypic assays

Isolates sequenced as part of the IPCD were donated by the Brown lab at Georgia Tech. We cultured isolates on LB agar plates, and confirmed their identity by sequencing the *lasR* gene, using colony *OneTaq* PCR (New England Biolabs). For protease activity, we inoculated colonies in 5 mL of LB broth, and grew strains in biological triplicates for 16-18 hours at 37 °C, 200 rpm. We pelleted cells and used the supernatant to run the Pierce Fluorescent Protease Assay Kit (Thermo). Readings were quantified using the TPCK Trypsin standard in triplicate, and normalized by strain optical density (OD at 600_nm_). All readings were compared to NPAO1. For colony autolysis experiments, we inoculated colonies in 5 mL of LB broth, and grew strains for 16-18 hours at 37 °C, 200 rpm. We tested colony autolysis by spotting 5 μL of overnight cultures on LB agar plates, and growing colonies for 24 hours at 37 °C.

## Author contributions

SPD and KO designed the study. KO, CYZ, and MM performed the *in silico* analysis of the data. KO performed the *in vitro* analysis. All authors contributed to the writing of the manuscript.

## Funding and acknowledgements

We wish to thank The Cystic Fibrosis Foundation (DIGGLE20G0) and The National Institute for Health (R01AI153116) to SPD for funding. We also thank members of the Diggle Lab and Jon Gerhart for helpful discussion. We thank the Brown Lab at Georgia Tech for donating strains from the IPCD collection.

---

**Table S1.** Nucleotide sequence mutation frequency

| Gene | Mutants | Mutation Frequency (%) |
| --- | --- | --- |
| *lasR* | 553 | 64.91 |
| *lasI* | 290 | 34.04 |
| *rhIR* | 849 | 99.65 |
| *rhII* | 847 | 99.41 |
| *pqsA* | 843 | 98.94 |
| *pqsB* | 685 | 80.40 |
| *pqsC* | 842 | 98.83 |
| *pqsD* | 843 | 98.94 |
| *pqsE* | 479 | 56.22 |
| *pqsH* | 845 | 99.18 |
| *pqsL* | 847 | 99.41 |
| *pqsR* | 789 | 92.61 |
| *qscR* | 772 | 90.61 |
| *rsaL* | 218 | 25.59 |
| *vqsR* | 559 | 65.61 |

| S1. Nucleotide sequence mutation frequency |  |  |
| --- | --- | --- |
| Gene | Mutants | Mutation Frequency (%) |
| lasR | 553 | 64.91 |
| lasI | 290 | 34.04 |
| rhIR | 849 | 99.65 |
| rhII | 847 | 99.41 |
| pqsA | 843 | 98.94 |
| pqsB | 685 | 80.40 |
| pqsC | 842 | 98.83 |
| pqsD | 843 | 98.94 |
| pqsE | 479 | 56.22 |
| pqsH | 845 | 99.18 |
| pqsL | 847 | 99.41 |
| pqsR | 789 | 92.61 |
| qscR | 772 | 90.61 |
| rsaL | 218 | 25.59 |
| vqsR | 559 | 65.61 |

S2: Mutation profile of QS genes for 12 isolates

**S2:**
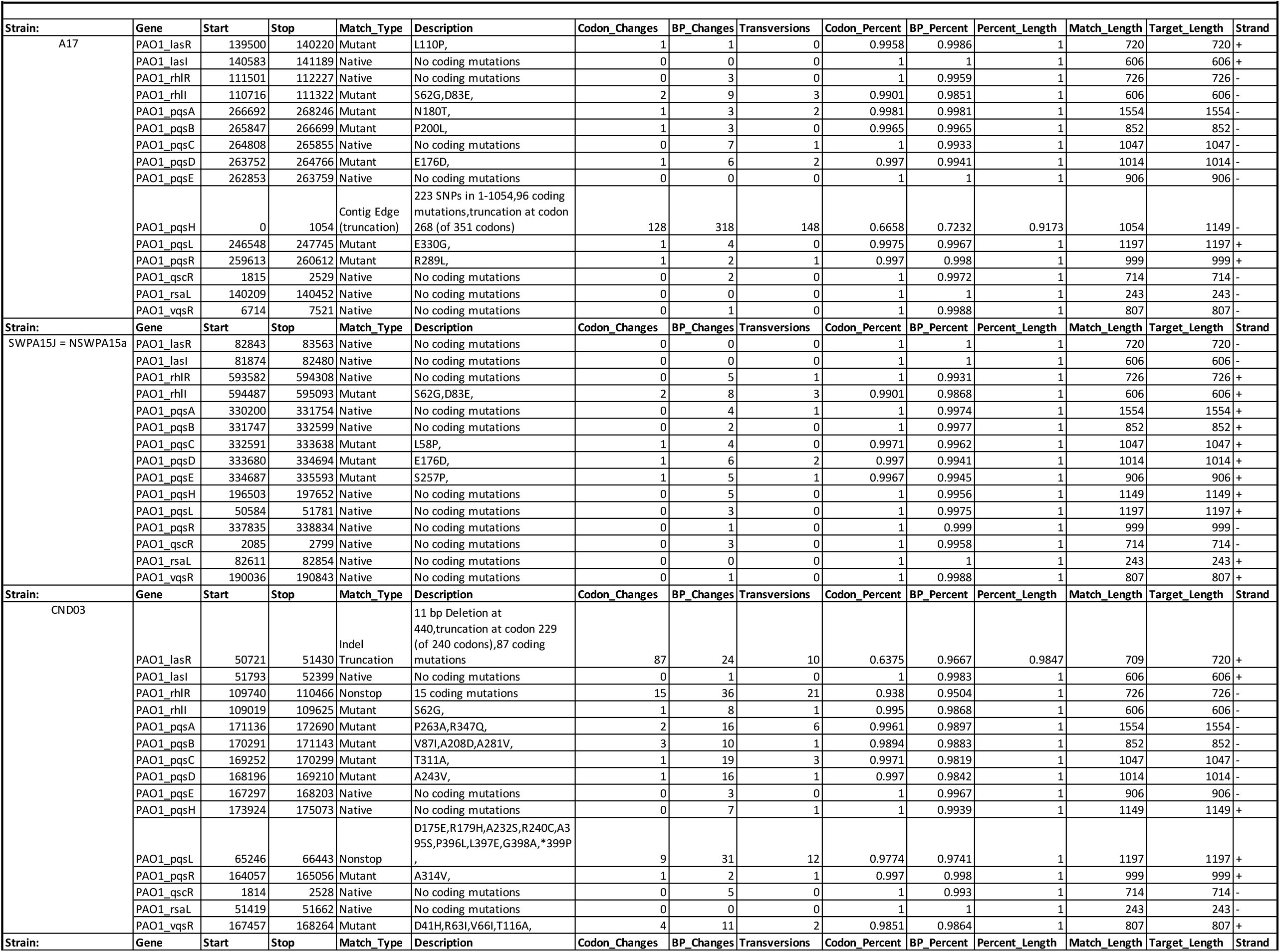
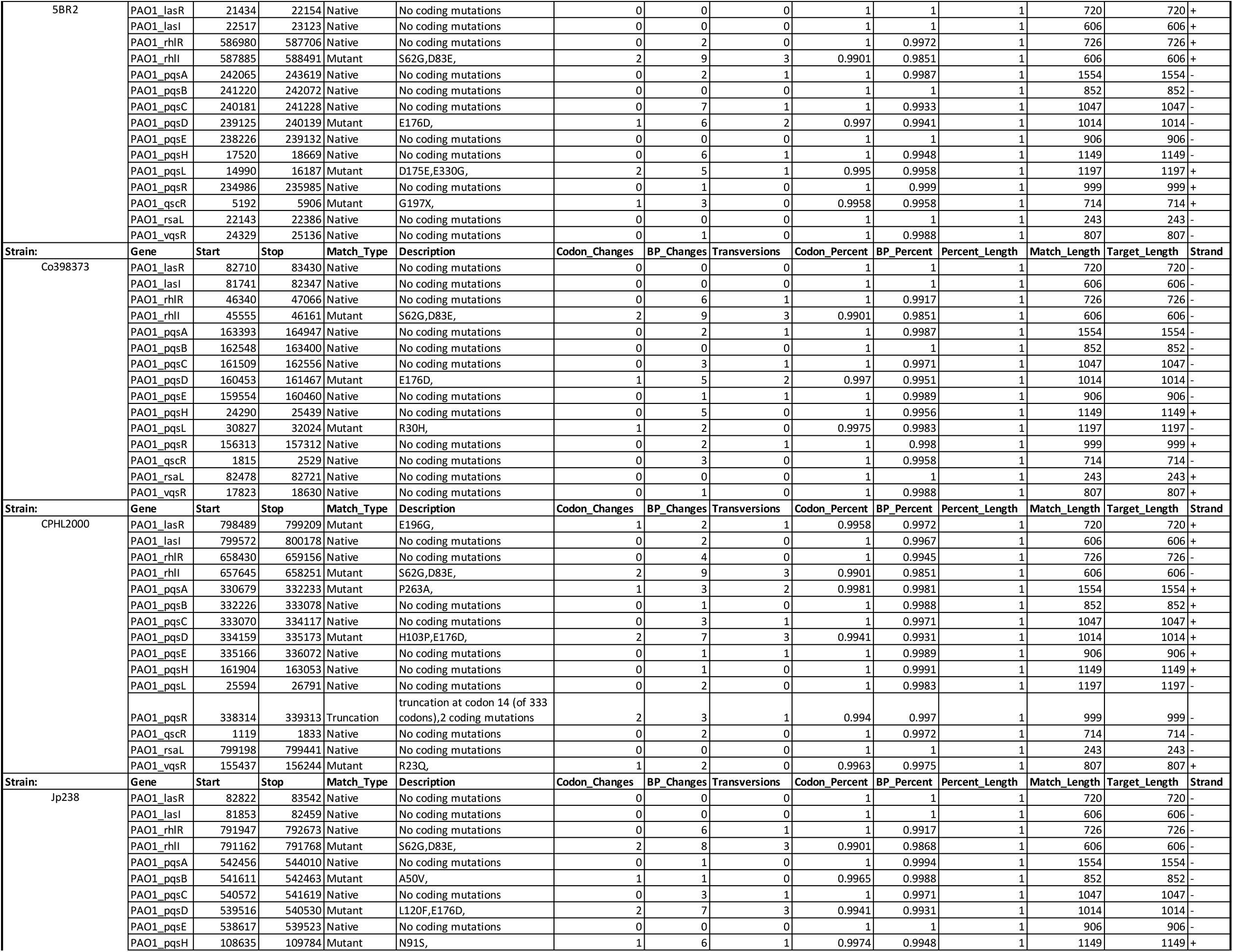
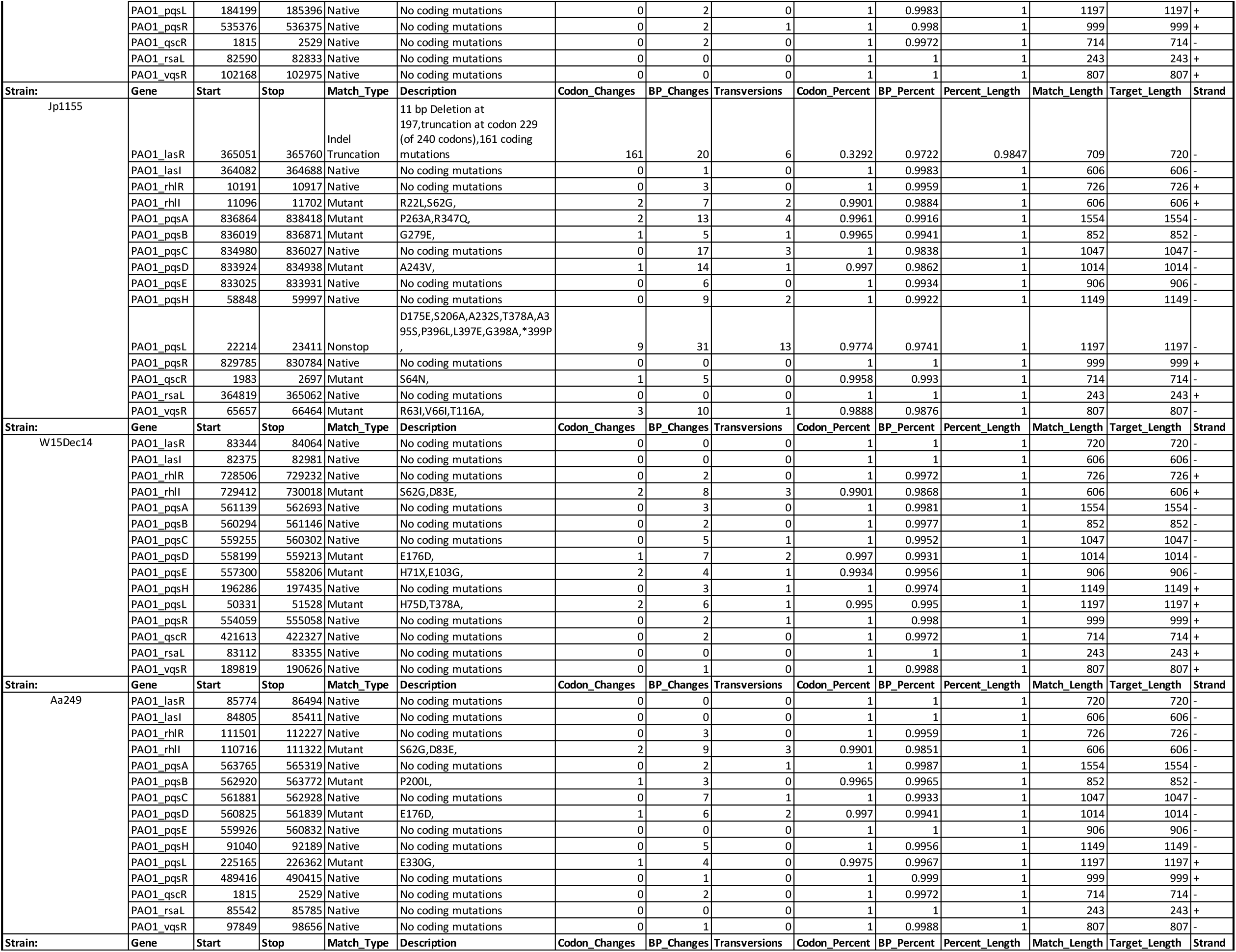
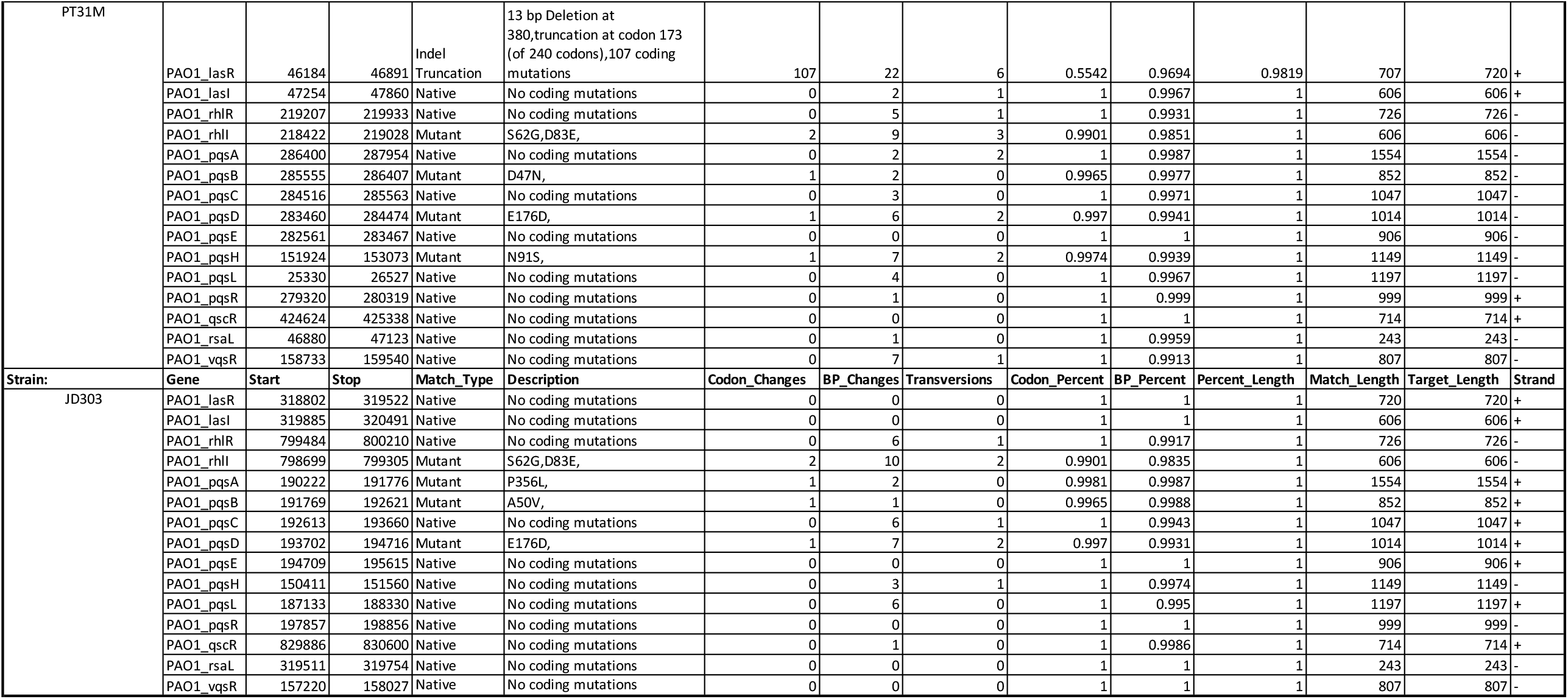
Mutation profile of QS genes for 12 isolates

